# GENERALIST: An efficient generative model for protein sequence families

**DOI:** 10.1101/2022.12.12.520114

**Authors:** Hoda Akl, Brooke Emison, Xiaochuan Zhao, Arup Mondal, Alberto Perez, Purushottam D. Dixit

## Abstract

Generative models of protein sequence families are an important tool in the repertoire of protein scientists and engineers alike. However, state-of-the-art generative approaches face inference, accuracy, and overfitting-related obstacles when modeling moderately sized to large proteins and/or protein families with low sequence coverage. To that end, we present a simple to learn, tunable, and accurate generative model, GENERALIST: *GENERAtive nonLInear tenSor-factorizaTion* for protein sequences. Compared to state-of-the-art methods, GENERALIST accurately captures several high order summary statistics of amino acid covariation. GENERALIST also predicts conservative local optimal sequences which are likely to fold in stable 3D structure. Importantly, unlike other methods, the density of sequences in GENERALIST-modeled sequence ensembles closely resembles the corresponding natural ensembles. GENERALIST will be an important tool to study protein sequence variability.

## Introduction

Advances in *omics* technologies allow us to investigate sequences of evolutionarily related proteins from several different organisms. Surprisingly, even when the function and structure are conserved, sequences within protein families can vary substantially^1^. This variability is governed by a combination of factors, including protein stability^2^, interaction partners^3^, and function^4^. Therefore, it is not feasible to rationalize observed variation in protein sequences using bottom-up mechanism driven models.

To understand the forces that constrain protein sequence variability and to identify new protein sequences that perform desired functions, we need methods to sample sequences that are likely to result in functional proteins^5^. Generative models of protein families that use multiple sequence alignments (MSAs) are one such approach. These models attempt to learn the covariation between amino acids across different positions and model a distribution over the sequence space that captures aspects of the observed covariation. The Potts model is one of the most popular generative models of protein families^6^. Potts model is a maximum entropy model constrained to reproduce positional amino acid frequencies and position-position pair correlations. Even though only 1- and 2-site frequencies are constrained, the model can reproduce higher order covariation statistics^7^. The model is easy to interpret, as it assigns an energy to sequences. In addition to modeling covariance between amino acid positions, Potts models have also been used to rationalize effects of mutations on fitness^8^, and to predict physical contacts between residues^7^.

However, there are significant issues with the Potts model. The associated numerical inference is computationally inefficient^9^, limiting their application to small proteins and protein domains (*L* ~ 100 residues). In comparison, median protein size in many organisms including humans is much larger (~ 350 residues)^10^. Due to the numerical inefficiencies in inference, there is no realistic way to tune the model beyond one- and two-position moments, for example, by incorporating multi-position correlations. Moreover, the model has many hyperparameters, including pseudocounts^11^ for unobserved amino acids and parameters related to phylogenetic reweighting^12^. How model predictions depend on these hyperparameters is not always clear. Finally, as we will show below, the Potts model does not reproduce statistics related to the density of sequences and result in highly unnatural optimal sequences. Field theoretic approaches^13^ can systematically generalize the Potts model by incorporating higher order epistasis. However, these models can only be trained on *very* small sequences. Another recent generalization that combines elements of autoregressive modeling and the Potts model; the autoregressive DCA model^14^, addresses the numerical issues associated with the Potts model. However, as we show below, this approach does not reproduce statistics related to the density of sequences and overfits the data when modeling families of large proteins with small MSAs.

Deep generative (DG) models are a potential alternative^15^ to Potts models for realistically sized proteins. However, DG models require large amounts of training data and lack interpretability. While sequencing advances have led to large MSAs, especially for bacterial protein families, many human proteins only exist in mammals and other higher order organisms where the MSA sizes are currently limited by the number of sequenced genomes and ultimately by the total number of mammalian species^16^. Neural network architectures are notorious for being over parametrized, including several hyperparameters for training the networks. Finally, as we show below, NN-based generative models may not necessarily improve in accuracy with the increasing complexity of the architecture.

Therefore, there is an urgent need for efficient, tunable, and accurate generative models. To that end, we present here GENERALIST: ***GENERA**tive non**LI**near ten**S**or-factoriza**T**ion*-based model for protein sequences and other categorical data. In GENERALIST, we model individual protein sequences in the data as arising from a sequence-specific Gibbs-Boltzmann distribution^17,18^. The energies of the distribution are shared across all sequences and the temperatures are assigned in a sequence-specific manner. The modeler only specifies complexity of the model (see below), and both the energies and the temperatures are inferred directly from the data. The temperatures embed individual sequences in a latent space which can be tuned to achieve a user-desired tradeoff between the novelty of generated sequences and the accuracy of the ensemble in reproducing properties of the natural MSA.

We use GENERALIST to model sequence variability in proteins that span multiple kingdoms of life, alignment sizes, and sequence lengths. We compare the performance of GENERALIST with three other generative models, the Potts model (referred to as adabmDCA^9^), the autoregressive DCA model (referred to as ArDCA^14^), and a variational autoencoder-based model (referred to as VAE^19^). We show that compared to these other models, GENERALIST captures higher order statistics of amino acid covariation across sequences. GENERALIST also predicts conservative local optima that are likely to fold in stable three-dimensional structures. Importantly, the ensemble of sequences generated using GENERALIST most accurately represents the density of sequences observed in nature. We believe that GENERALIST will be an important tool to model protein sequences and other categorical data.

## Results

### The Mathematical formalism of GENERALIST

In GENERALIST (Figure 1), we start with a one-hot encoded representation of a multiple sequence alignment of *N* sequences of length *L*; *σ_nla_* = 1 if the amino acid at position *l* in the protein sequence indexed *n* has the identity *a*. Sequences are modeled as arising from their own Gibbs-Boltzmann distribution^17,18^:

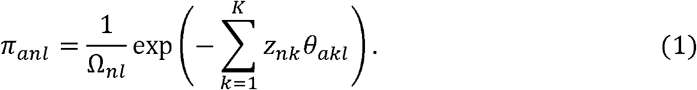

**Figure 1.**
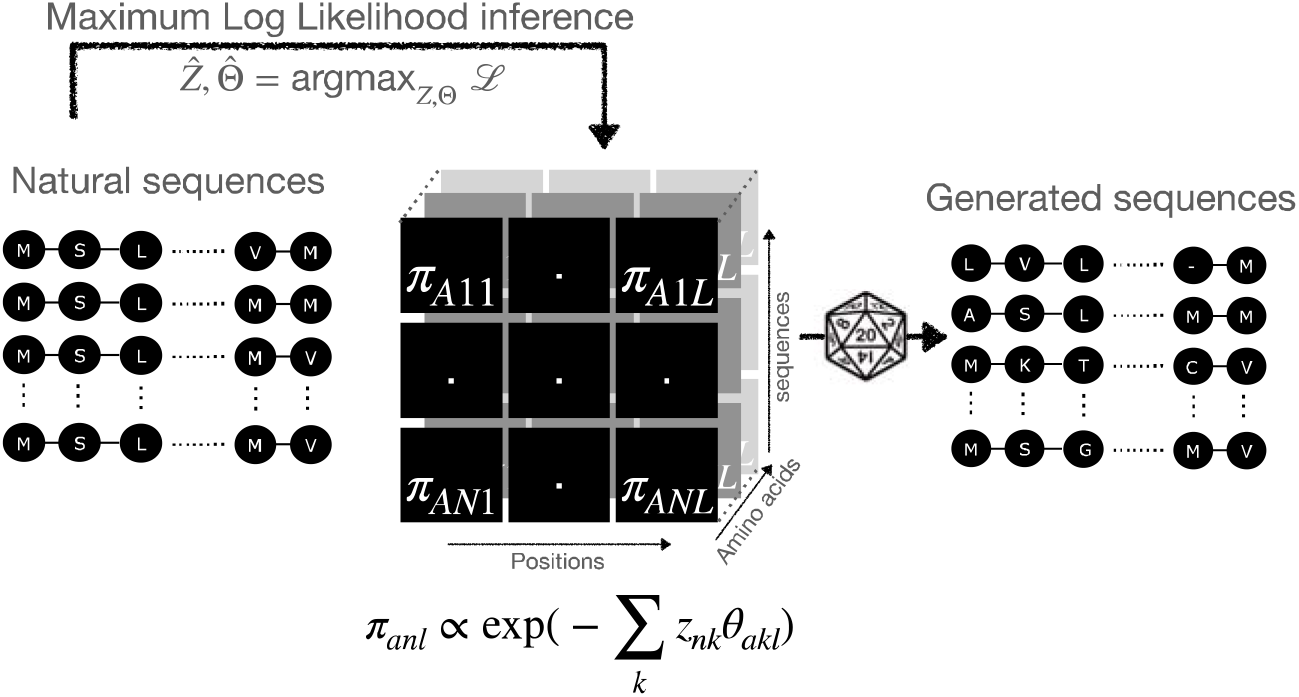
Schematic of the GENERALIST approach. Sequences are modeled as arising from their own Gibbs-Boltzmann distributions over categorical variables. The inferred probabilities are used to generate new sequences.

In Eq. (1), *z_nk_* are sequence-specific inverse temperature-like quantities (latent space embeddings), *θ_akl_* are position and amino acid dependent variables, and Ω_nl_ is the partition function that normalizes the probabilities. We can write down the total log-likelihood of observing the data:

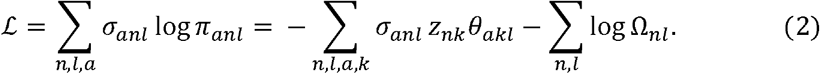

The gradients of the log likelihood with respect to position- and amino-acid dependent parameters *θ_akl_* and *z_nk_* are analytical. The parameters are simultaneously inferred using maximum likelihood inference. Once the parameters are inferred, sequences can be sampled in the vicinity of any sequence in the MSA using probabilities inferred in Eq. (1).

Below, we present our results for two proteins: Bovine Pancreatic Trypsin Inhibitor or BPT1, a small protein domain comprising ~ 50 amino acids with a large MSA of ~ 16000 sequences and epidermal growth factor receptor or EGFR, a large protein comprising ~ 1000 amino acids with a small MSA of ~ 1000 sequences. In the SI, we show our analyses for dihydrofolate reductase or DHFR (~ 160 amino acids, ~ 7000 sequences in the MSA), p53 (~ 350 amino acids, ~ 800 sequences in the MSA), and mammalian target of rapamycin or mTor (~ 2500 amino acids, ~ 500 sequences in the MSA). Details of model training can be found in SI Section 1.

### Choosing the optimal latent space dimension in GENERALIST

GENERALIST is a latent space model. Increasing latent space dimension typically improves the ability of the generated ensemble to accurately capture summary statistics of the data (for example, amino acid frequencies and covariation). At the same time, a high dimensional latent space can result in a generated ensemble that is nearly identical to the natural one; trivially reproducing all statistics but failing to generate new sequences. Therefore, a common challenge with latent space models is selecting an appropriate dimension to avoid overfitting.

GENERALIST offers a natural way of evaluating overfitting. We computed for each generated sequence the fractional Hamming distance (fraction of positions that have a different amino acid) to the closest natural sequence (blue distributions in Figure 2, SI Section 2). We compared these distributions to the distribution of nearest neighbor distances within the natural ensembles (gray distributions in Figure 2). For an overfit ensemble, the distribution will peak sharply at zero; implying that generated sequences are nearly identical to natural ones. In Figure 2, we show these distance distributions for GENERALIST ensembles trained with different latent space dimensions. When the latent space dimension is low, GENERALIST ensembles comprise sequences that are on average different from the natural sequences (as quantified by the mean fractional Hamming distance to the closest natural sequence, blue bar). However, the ensembles tend towards overfitting with higher latent space dimension, as seen in the leftward shift in the distribution of distances to the nearest natural neighbor.

**Figure 2.**
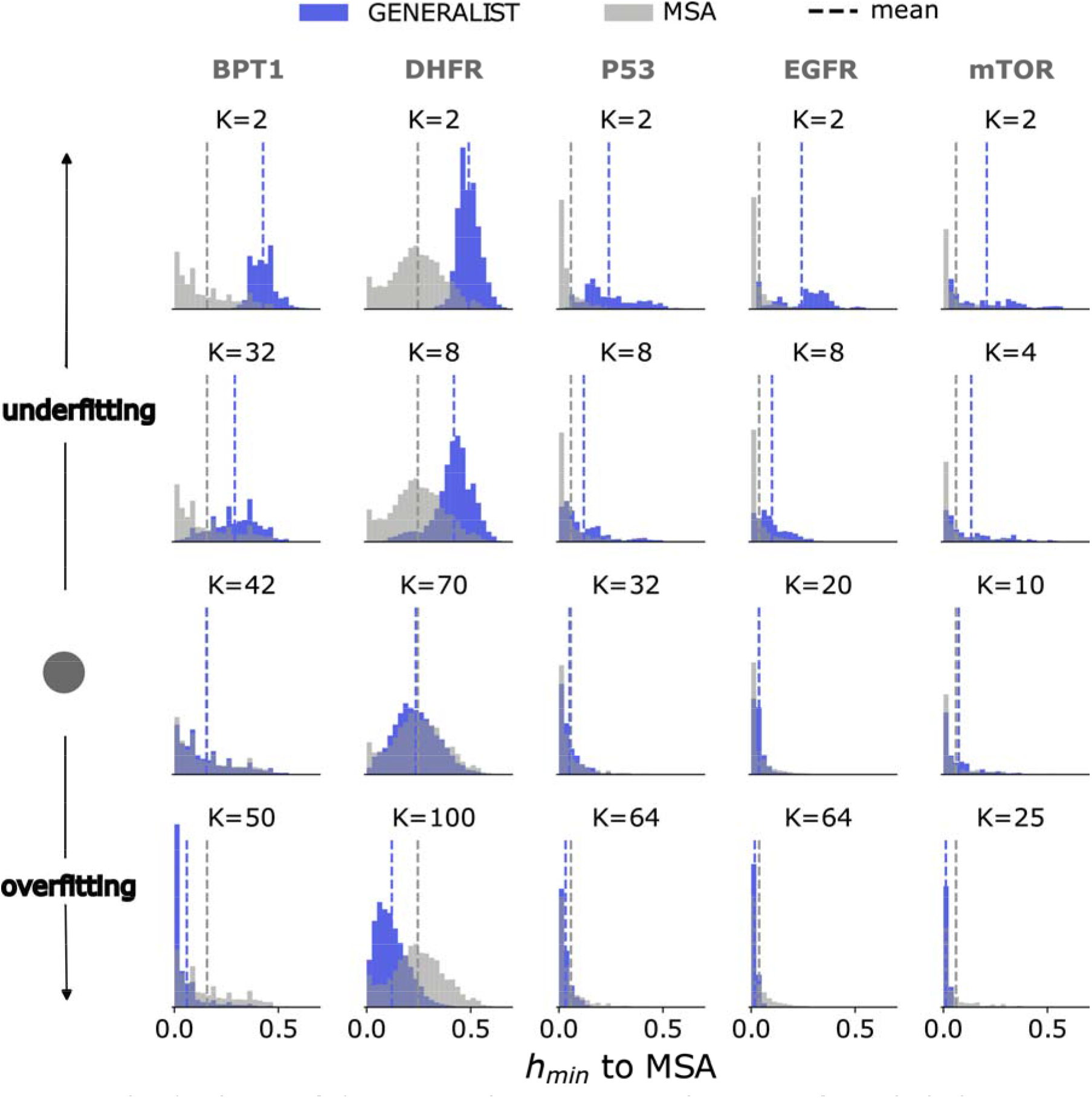
The distribution of distances to the nearest natural sequence for multiple latent space dimensions. For each protein and a given latent space dimension, an *in silico* ensemble was generated using GENERALIST. For each generated sequence, the minimum fractional Hamming distance to the natural ensemble was evaluated (blue). The same calculation was repeated for natural sequences (gray). The dashed vertical lines represent the means of the distributions. The gray disc on the left indicates the optimal latent space dimension for each protein.

In the middle, we find the optimal latent space dimension as the one that matches the average separation between nearest neighbors in natural sequences (dashed gray line) and the average separation between sequences in the generated ensemble and the nearest natural neighbor (dashed blue line). For the rest of the analyses, we choose this optimal dimension for the studied proteins. Notably, variational autoencoders lend themselves to a latent space description as well. Yet, we observed that ensembles generated using VAEs did not exhibit a systematic trend towards overfitting when the latent space dimension was increased (SI Figure 1).

### GENERALIST reproduces high order summary statistics of natural sequences

A key metric to evaluate the accuracy of generative models is their ability to reproduce summary statistics on the sequences (SI Section 3). In Figures 3A and 3B, we show for BPT1 and EGFR that GENERALIST accurately reproduces amino acid frequencies and mean removed positional correlations up to order 4. Notably, as seen in Figure 3C and 3D (SI Figure 2), while adabmDCA, ArDCA, and VAE-based predictions of positional frequency statistics correlate strongly with those observed in the natural sequences (SI Section 4); these methods typically under-predict these statistics (quantified by the slope of the best fit line).

**Figure 3.**
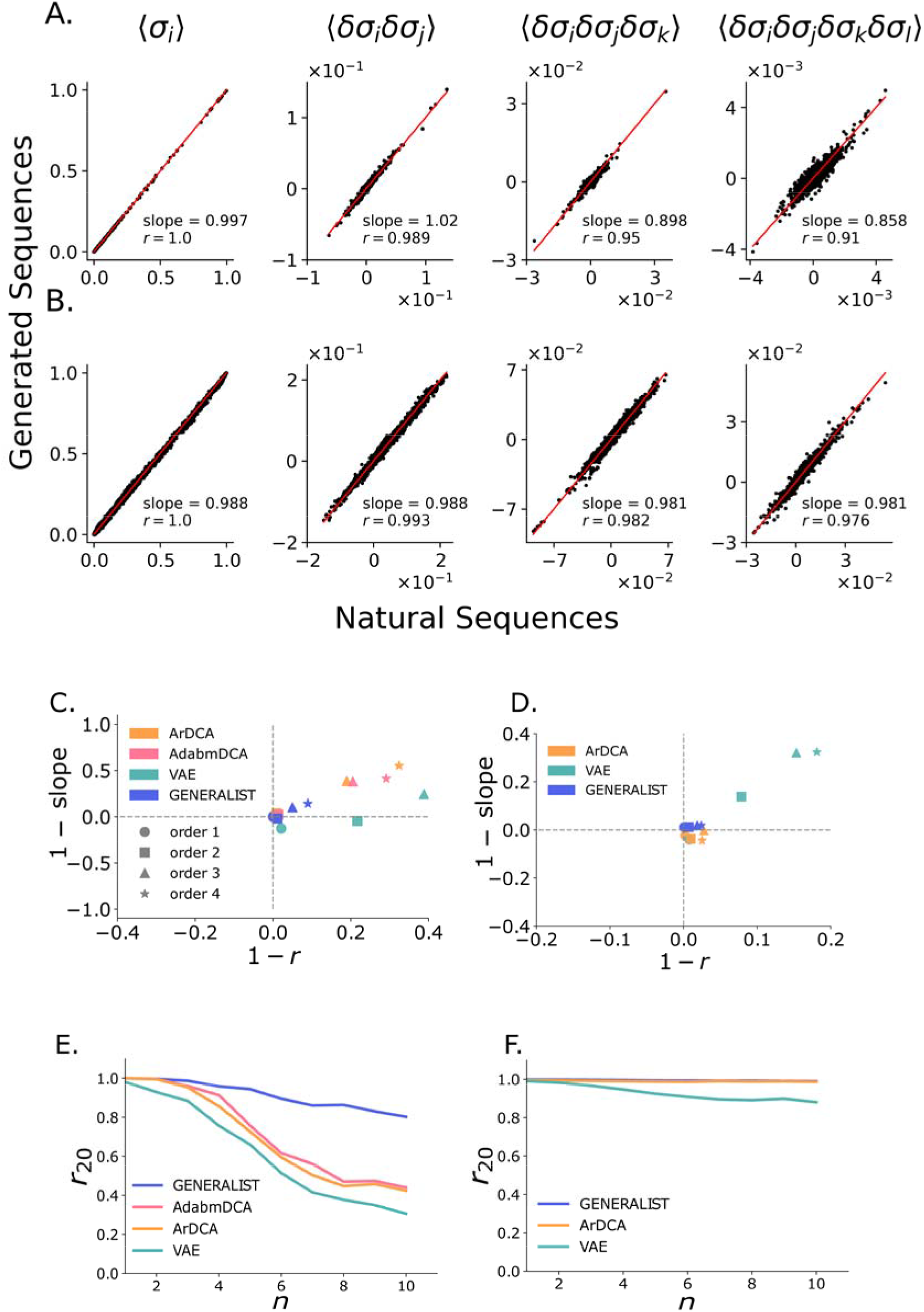
**Panels A and B.** Comparison of amino acid frequencies, mean removed pair, three and four body correlations calculated from GENERALIST-generated in silico ensembles (y-axis) and the natural sequences (x-axis) for BPT1 (panel A) and EGFR (panel B). **Panels C and D.** 1 – Pearson correlation coefficient versus 1 – slope of the best fit line for the comparison between amino acid frequencies, mean removed pair, three and four body correlations for GENERALIST, ArDCA, adabmDCA, and VAEs shown for BPT1 (panel **C)** and EGFR (panel **D). Panels E and F.** The average Pearson correlation coefficient between frequencies of top 20 amino acid combinations of order n (x-axis) averaged across different combinations (y-axis) for GENERALIST, ArDCA, adabmDCA, and VAEs shown for BPT1 (panel **E)** and EGFR (panel **F**).

Next, we investigated the ability of the generated ensembles to reproduce very high order summary statistics. Most amino acid combinations of order higher than 4 are rarely found in natural MSAs. We therefore used a recently introduced metric *r*_20_ that measures the average Pearson correlation between the occurrence frequency of the top 20 amino acid combinations of any given order^20^. In Figure 3E and 3F (SI Figure 3), we show that GENERALIST accurately captures co-occurrence frequencies of the most frequent amino acid combinations up to order 10. The ability of GENERALIST to capture these higher order statistics did not depend on restricting our attention to the top 20 amino acid combinations (SI Figure 4). In comparison, adabmDCA, ArDCA, and VAEs led to less accurate predictions about higher order correlations when the MSAs were large (BPT1 in the main text and DHFR in the SI). Importantly, the ensembles generated using VAEs did not exhibit a systematic trend toward more accurate predictions when the latent space dimension was increased (SI Figure 1). Finally, ArDCA could capture higher order positional correlations for large proteins with small MSAs (Figure 3F, SI Figure 3). However, as we will show below, this was due to overfitting.

These results conclusively show that GENERALIST-based sequence ensembles retain positional correlation information of arbitrarily high orders observed in naturally occurring sequences for large proteins as well as for proteins with very small MSAs.

### GENERALIST finds conservative optimal sequences

A key feature of generative models is the ability to assign probabilities to arbitrary sequences and therefore find local sequence optima (sequences corresponding to the local maximum of the probability). The local optima inform us about the local structure of the inferred sequence space energy landscape and their relationship to naturally occurring sequences. For example, if the generative models are purely data-driven, that is, if they do not incorporate any information about structure/function/fitness, it may be desirable that the local optima are in the vicinity of natural sequences.

To test the relationship between local optimum sequences and natural sequences, we use GENERALIST, adabmDCA, and ArDCA to obtain locally optimal sequences. VAE was not included because VAEs involve a nonlinear transformation from the latent space to the sequence space and therefore the probability in the sequence space is difficult to calculate.

We obtained local minima in adabmDCA and ArDCA using a random search (SI Section 5). Briefly, we start from sequences in the natural MSA and randomly mutated amino acids while only accepting mutations that improve sequence probability as evaluated by the model. Multiple iterations of this operation lead to local optimum sequences. The local optimum sequences predicted by GENERALIST were obtained by finding the highest probability sequence corresponding to the latent space embedding of natural sequences. This analysis was only performed on BPT1 where all three models could be trained in a reasonable time.

As seen in Figure 4A, adabmDCA generates locally optimal sequences that differed by a staggering 84% from the closest naturally occurring sequence neighbor. These optimal sequences were predicted to be significantly better compared to the starting natural sequences, with an average improvement by ~ 110 fold in probability at each position (with a total average increase in probability by a factor of ~ 5 × 10^104^ when considering the entire sequence) (Figure 4B, measured by log odds ratio). These local minima in the Potts model that do not resemble any natural sequences are reminiscent of the unwanted spurious minima in Hopfield networks^21^. Compared to adabmDCA, ArDCA generated local optimal sequences that were significantly more conservative (on an average, 17% difference compared to 84%) (Figure 4A). The optimal sequences were also predicted to be a relatively modest improvement over the starting natural sequence with an improvement by ~ 1.5 fold in probability at each position with a total average increase in probability by a factor of ~ 5 × 10^9^ when considering the entire sequence (Figure 4B). Like ArDCA, GENERALIST-based local optima were significantly more conservative. As seen in Figure 4A, the local optimum sequences differed from the closest naturally occurring sequences on an average by 8%. As seen in Figure 4B, the per amino acid improvement was only ~ 1.1 fold with a total average increase in probability by a factor of ~ 7 × 10^2^ when considering the entire sequence.

**Figure 4.**
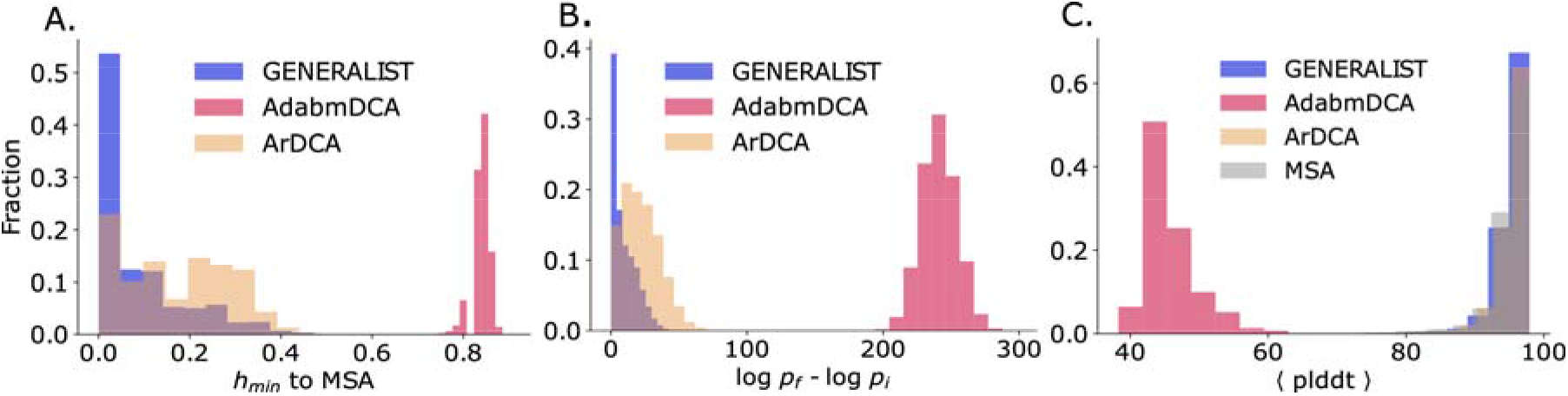
**Panel A.** The distribution of distances to the nearest natural neighbor from sequences optimized using GENERALIST, ArDCA, and adabmDCA modeled probabilities. **Panel B.** The log-fold improvement in probabilities between the starting sequence and the local optimum. **Panel C.** Sequence-averaged plddt scores for AlphaFold2 predicted structures for the locally optimum sequences.

To test whether these sequences potentially fold in stable 3D structures, we used AlphaFold2^22^, a recent machine learning method that can predict 3D structures from sequences and MSAs (SI Section 6). We used the sequence-averaged predicted local distance difference test (plddt) as a proxy for quality of predicted structures. Previous studies have shown that a sequence average plddt of > 80 corresponds to sequences that are likely to fold in stable 3D structures^23^. As seen in Figure 4C, local optimal sequences imputed by adabmDCA were predicted to be significantly worse folders compared to both GENERALIST and ArDCA. While ArDCA and GENERALIST produce sequences that were predicted to be comparable by AlphaFold2 on average.

These results show that GENERALIST can identify local optima in the sequence space. These optima are typically not seen in nature. The optimal sequences predicted by GENERALIST were also predicted by AlphaFold2 to fold in stable 3D structures

### GENERALIST reproduces statistics related to the density of sequences in the natural ensemble

In addition to reproducing the summary statistics (Figure 3), an important test for generative models is capturing the density of sequences in the natural ensemble^7,24^. To that end, we evaluated three different statistics for all generated ensembles: (a) the distribution of distances between pairs of randomly picked sequences, (b) the distribution of nearest neighbor distances, and (c) the distribution of distances to the nearest natural neighbor.

In Figure 5A and 5B (SI Figure 5), we plot the distribution of fractional Hamming distances between pairs of random sequences in an ensemble. We see that all generative models, except for the VAEs, accurately reproduced this distribution, implying that most ensembles captured the expanse of the natural sequence ensemble. Ensembles generated using VAEs comprised sequences that are on average are closer to each other than natural sequences.

**Figure 5.**
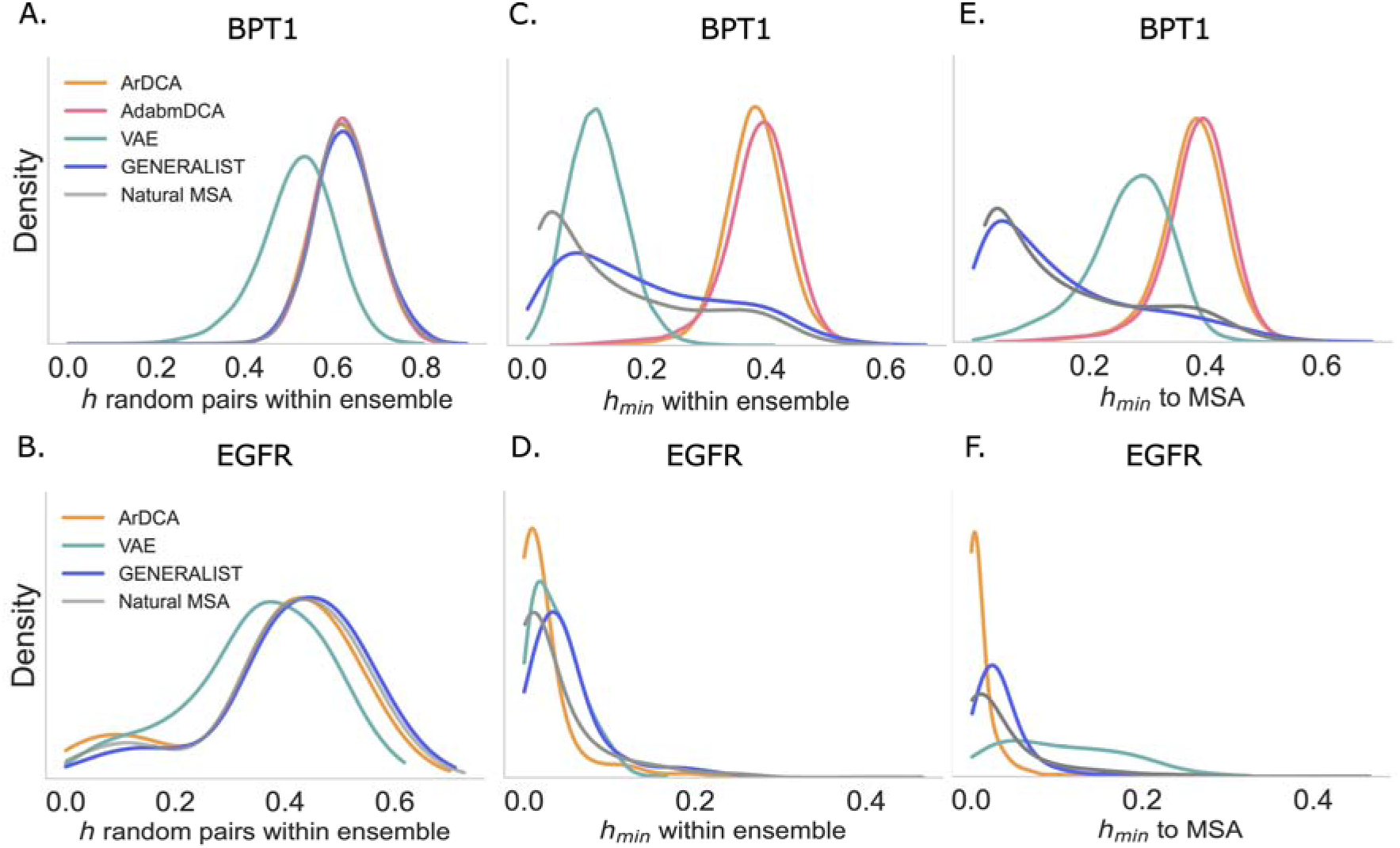
**Panels A and B.** Distribution of fractional Hamming distances between random pairs of sequences within an ensemble shown for BPT1 (panel A) and EGFR (panel B). **Panels C and D.** Distribution of fractional Hamming distances to the closest sequence within an ensemble for different models shown for BPT1 (panel C) and EGFR (panel D). **Panels E and F.** Distribution of fractional Hamming distances to closest natural sequence for different models shown for BPT1 (panel E) and EGFR (panel F).

The distribution of nearest neighbor distances portrayed a more complex picture. In Figure 5C and 5D (SI Figure 6, SI Section 2), we plot the distribution of fractional Hamming distances to the nearest neighbor within the ensemble. When MSAs were large (BPT1 in Figure 5C and DHFR in SI Figure 6), ArDCA/adabmDCA generated ensemble comprised sequences that were farther away from each other compared to natural sequences. In contrast, for small MSAs (EGFR in Figure 5D and p53 and mTor in SI Figure 6), ArDCA generated ensembles comprised sequences that were closer to each other compared to natural sequences. VAEs generated ensemble always comprised sequences that were on average closer to each other than natural sequences (with the exception of mTor). In contrast GENERALIST generated ensemble closely reproduced the density of nearest neighbor sequences observed in the natural ensembles.

Next, we compared the distance distribution to the nearest natural neighbor. Here too, GENERALIST generated ensembles closely reproduced the density of nearest neighbor sequences (Figure 5E and 5F, SI Figure 7). In contrast, ArDCA/adabmDCA generated sequences were farther from the natural sequences if the MSA was large (BPT1 in Figure 5E and DHFR in SI Figure 7). ArDCA generated ensembles for proteins with small MSAs were overfit to the MSA as evidenced by a distribution of distances with a sharp peak at zero (EGFR in Figure 5F and p53 and mTor in SI Figure 7). This overfitting also explains the accuracy with which ArDCA can reproduce sequence summary statistics for EGFR and other large proteins with small MSAs (Figure 3F, SI Figure 3). Finally, VAEs generated ensemble comprised sequences that were farther away compared to natural sequences for all proteins.

These results show that an optimally tuned GENERALIST ensemble can capture various aspects of the density of sequences in the natural ensemble.

## Discussion

Generative models of protein sequence families are an important tool for protein scientists and engineers alike. Ideally, these models should be simple to learn, tunable, and accurate, especially when studying proteins of significant clinical interest which tend to be large proteins with small MSAs.

In this work, we examined three state-of-the-art models. Physics-based Potts models could only be used to model small sequences, limiting their application to single domains and small proteins. Moreover, these models could not be tuned. The sequence ensemble generated by Potts models could not reproduce the density of sequences in the natural ensemble and had optima that appeared unnatural. In contrast, the autoregressive generalization of the Potts model was significantly more efficient in model fitting and more accurate in reproducing summary statistics such as frequencies of higher order amino acid combinations (Figure 3). The model also reproduced reasonable local optima that were computationally deemed to fold in stable 3D structures (Figure 4). However, the autoregressive Potts model could not reproduce the density of sequences in the natural ensemble (Figure 5). Importantly, given that the model has *O*(*L*^2^) parameters for proteins of sequence length *L*, this model overfits the training data when modeling human proteins of significant clinical interest which are large and have small MSAs.

Neural networks based variational autoencoders were efficient and did not appear to overfit the training data (Figure 5). Overall, these models were less accurate in predicting summary statistics of sequences compared to GENERALIST, the Potts model, and the autoregressive generational of the Potts model. At the same time, potentially owing to model complexity (and therefore parameter non-identifiability) and low amounts of training data, the models appeared to not have any systematic trends with respect to accuracy and overfit as a function of the dimension of the latent space.

In contrast, GENERALIST is efficient, accurate, and tunable, allowing us to analyze large proteins with small MSAs. Notably, given its simple structure, there are several avenues of improving GENERALIST. For example, function/fitness information obtained from deep mutational scanning^15^ can be incorporated as constraints on the energies and phylogenetic information can be imposed as constraints on the latent space. Finally, GENERALIST can be easily reformulated for any other categorical data, for example, presence/absence of single nucleotide polymorphisms or nucleotide sequences. We believe that GENERALIST will be an asset for protein scientists and engineers alike.

## Acknowledgments

HA, BE, and PD are supported by NIGMS grant R35GM142547. The authors would like to thank Francesco Zamponi and Martin Weigt for fruitful discussions about implementation of the Potts model and the autoregressive Potts model. We would like to thank Juannan Zhou for critical comments and useful discussions.

