## Supplementary Information for "GENERALIST: An efficient generative model for protein sequence families"

Hoda Akl[1*], Brooke Emison[1], Xiaochuan Zhao[1], Arup Mondal[2], Alberto Perez[2], and Purushottam Dixit[1,3,4,5*]

[1]1Department of Physics, University of Florida, Gainesville, FL 32603

[2]Department of Chemistry, University of Florida, Gainesville, FL 32611

[3]Department of Chemical Engineering, University of Florida, Gainesville, FL 32603

[4]UF Genetics Institute, University of Florida, Gainesville, FL 32608

[5]UF Health Cancer Center, University of Florida, Gainesville, FL 32610

[*]Purushottam Dixit: , Hoda Akl:

All scripts can be found on: https://github.com/hodaakl/GENERALIST

#### 1 Maximum log likelihood inference

We consider a multiple sequence alignment (MSA) of $N$ sequences where each position could be one of $D = 21$ categories (20 amino acids + alignment gap). The MSA can be represented by a 3-D binary object $\sigma_{anl}$ where $a \,\epsilon\, \{1..D\}$ , $n \,\epsilon\, \{1..N\}$, $l \,\epsilon\, \{1..L\}$, $\sigma_{anl} = 1$ *if* position $l$ in sample $n$ is occupied by amino acid/category $a$, $\sigma_{anl} = 0$ otherwise. Each sequence is modeled as sampled from a Gibbs-Boltzmann distribution parameterized by sequence-specific "inverse temperature" variables $\vec{z}_n$ and "energies" $\vec{\theta}_{al}$ that are shared between the sequences. These parameters have dimension $K$ that is user-specified. In this model, the probability that position $l$ in sequence $n$ is occupied by category (amino acid or a gap) $a$ ($\sigma_{anl} = 1$) is given by

$$\pi_{anl} = \frac{1}{\Omega_{nl}} \exp\left(-\sum_{k=1}^{K} z_{nk}\theta_{akl}\right). \tag{1}$$

In Eq. 1, $\Omega_{nl} = \sum_d \exp(-\sum_k z_{nk}\theta_{dkl})$ is a normalization constant.

#### 1.1 Maximizing Log Likelihood

The likelihood of observing an amino acid sequence $\sigma_n$ is $P(\sigma_n|z_n,\theta) = \prod_{a,l} \pi_{anl}^{\sigma_{anl}}$. The log likelihood of the entire MSA is therefore $\mathcal{L} = \log(\prod_{n,a,l} \pi_{anl}^{\sigma_{anl}})$, using the element-wise probability defined in Eq. 1, the log likelihood is

$$\mathcal{L} = -\sum_{l,n,a,k} \sigma_{anl} z_{nk} \theta_{akl} - \sum_{n,l} \log \Omega_{nl} \tag{2}$$

Maximizing the log likelihood allows us to arrive at the model parameters $z$s and $\theta$s. The derivatives of the log likelihood with respect to these parameters are given by

$$\begin{aligned}
\frac{\partial \mathcal{L}}{\partial z_{nk}} &= -\sum_{l,a} \sigma_{anl}\theta_{akl} + \sum_{l,a} \sigma_{anl} \frac{\sum_{d=1}^{D} \theta_{dkl} \exp\left(-\sum_{k=1}^{K} z_{nk}\theta_{dkl}\right)}{\Omega_{nl}} \\
&= -\sum_{l,a}^{L,D} \sigma_{anl}\theta_{akl} + \sum_{l,a,d} \sigma_{anl}\theta_{dkl}\pi_{dkl} \\
&= \sum_{l,a}^{L,D} \theta_{akl} \left(\pi_{anl} - \sigma_{anl}\right)
\end{aligned} \tag{3}$$

$$\begin{aligned}
\frac{\partial \mathcal{L}}{\partial \theta_{akl}} &= -\sum_{n} \sigma_{anl} z_{nk} + \sum_{n,d} \sigma_{dnl} z_{nk} \pi_{anl} \\
&= \sum_{n}^{N} z_{nk} \left(\pi_{anl} - \sigma_{anl}\right)
\end{aligned} \tag{4}$$

#### 1.2 Training

Elements of $z$ and $\theta$ are initialized from a uniform random distribution $[0,1]$ then rescaled through $z \leftarrow z/|z|$ and $\theta \leftarrow \theta/|\theta|$. This rescaling avoids numerical overflow issues. The optimization of $z$s and $\theta$s to maximize the log likelihood is done in an adaptive manner using ADAM optimization algorithm [1]. The parameters for ADAM are as follows: The exponential decay rate for the first moment estimates is 0.8. The exponential decay rate for the second-moment estimates is 0.999, the step size is 0.1 and finally, epsilon, which is a very small number to prevent division by zero, is $10^{-8}$. The stopping criteria for training is that $\frac{|\partial\mathcal{L}/\partial z|}{|z|} < 1$ and $\frac{|\partial\mathcal{L}/\partial\theta|}{|\theta|} < 1$. For each protein, we train GENERALIST for many latent dimensions spanning the range 2 to 100. We use the minimum Hamming distance to MSA distributions to decide the optimal latent dimension (see SI section 2 and main text Fig 2). The chosen latent dimensions are 42, 70, 32, 20 and 10 for proteins BPT1, DHFR, P53, EGFR and mTOR respectively. Once the optimal latent dimension is determined, it is used for all further analyses.

##### 1.3 Sampling from the model

The latent variables $z$ along with the energy-like parameters $\theta$ specify the inferred Gibbs-Boltzmann distributions from which samples are drawn. Each sample $n$ is represented by $\vec{z}_n$ in the latent space of dimension $K$. Those vectors can be used to generate sequences. After having learned the $z$s and $\theta$s, we sample $z$ with replacement $N$ times where $N$ is equal to the number of natural sequences in the MSA and use the sampled $z$s along with $\theta$s to probabilistically generate sequences according to Eq. 1.

##### 1.4 Data processing

We use five different protein families to test the different generative models: BPT1 (UniProt: P00974) domain position 40 - 90, DHFR (Pfam: PF00186), P53 (Pfam: PF00870), EGFR (UniProt: Q504U8), and mTOR (UniProt: P42345). We obtain the MSA for DHFR through Pfam database [2] and construct the MSA for the other protein families using Jackhammer [3].

We impose a similarity threshold of 30%, which sets a lower bound on the fractional Hamming distance between members of the protein family and the reference sequence used as the search seed to obtain the MSA. We only retain unique sequences in the MSA. We end up with BPT1 MSA of 16569 and 51 positions, DHFR MSA of 7164 sequences and 158 positions, P53 MSA of 785 sequences and 341 positions, EGFR MSA of 1010 and 1091 positions, and mTOR MSA of 529 sequences and 2549 positions. All MSAs are available on GitHub.

#### 2 Density of the Sequence Space

To measure the divergence between two sequences we calculate the fractional Hamming distance which is the fraction of positions in which two sequences vary.

We calculate the fractional Hamming distance between every generated sequence and all natural sequences. The natural sequence corresponding to the minimum fractional Hamming distance $h_{min}$ represents the closest natural neighbor to a given sequence. The values $h_{min}$ for all generated sequences define the distribution of distances to the closest natural neighbors. We estimate the nearest neighbor density within an ensemble, generated or natural, by calculating the fractional Hamming distance between every member of that ensemble and all the other members, again using $h_{min}$ to define the distribution of distance to the closest sequences within the same ensemble. Finally, we also calculate the fractional Hamming distance $h$ between random pairs within an ensemble, and we randomly generate 1000 pairs to get that distribution.

### 3 Statistical accuracy of generated ensembles

#### 3.1 Mean removed statistics

To assess the models' accuracy, we measure their ability to reproduce the frequency of amino acid combinations of different lengths. The simplest one is the $1^{st}$ order statistics, which is single site frequency. Here we have $21L$ frequencies (each amino acid -or gap- at every position). For statistics of order $n$ where $n \in [2..4]$, we randomly pick $n$ positions and obtain the amino-acid combination/"word" from a random sample in the natural MSA, then calculate its mean removed frequency in the natural and the generated ensemble, this is repeated for 7000 times. The Pearson correlation coefficient and the slope of the best fit line are evaluated between frequencies obtained from natural sequences and those obtained from generated ensembles.

#### 3.2 Calculating $r_m$

We used a previously published metric, $r_{20}$ [4], to assess the models' fidelity in reproducing frequencies of higher-order amino acid combinations. As described in [4] for any order $n$, we randomly picked $n$ positions and obtain all the unique "words"/amino-acid combinations that exist in that position set in the natural MSA. We use only the most 20 frequent words to compare with the generated dataset and obtain the Pearson correlation $r$. For each $n$ we use 100 position sets and the value $r_{20}$ is defined as the average of the Pearson correlation coefficients over the different position sets. For $r_{10}$ and $r_{50}$ we follow the same procedure using the most frequent 10 and 50 combinations respectively.

### 4 Benchmarking

We compare GENERALIST to three state-of-the-art generative models; ArDCA [5] , adabmDCA [6] and MSA-VAE [7]. We utilized the code published by the authors to train the different models. We use the default parameters suggested in the original publication unless otherwise stated. ArDCA as well as adabmDCA use weights for the natural sequences to capture the correlations that arise due to population structure, for both models we modify the function arguments to set the weights to be equal across sequences.

**ArDCA**: To specify equal weights we set the corresponding original code argument `theta = 0`. Since the MSA is preprocessed we do not require ArDCA function to further alter the MSA by setting the argument `max_gap_fraction = 1`, which ensures that the algorithm does not remove any of the sequences present in the MSA due to a threshold set on the gap fraction of the sequence. We used $L2$ regularization strength of $\lambda_J = 10^{-4}$ and $\lambda_h = 10^{-6}$ as recommended for generative tests in the original publication. The ArDCA model optimizes the probability of the sequences in the Entropic order by default, from least to most variable positions. This choice of ordering informs our calculation of the sequence probability using ArDCA. After ArDCA is trained we extract the following parameters: single site fields `H`, two site couplings `J`, the vector corresponding to the how the positions are permuted `idxperm`, and the probability of the initial site in the sequence given the chosen ordering `p0`. We use those parameters to calculate the probability of a sequence according to the ArDCA model for the local minima analysis. Code for ArDCA probability calculation using the model parameters is available on github.com/hodaakl/GENERALIST/benchmark/ArDCA.

**adabmDCA**: Similar to ArDCA, we used equal weights for all sequences by setting the sequence similarity threshold parameter to zero. A pseudocount was used to account for unobserved amino acids/amino acid pairs, set to the default value of $1/M_{\text{eff}}$ where $M_{\text{eff}}$ is the effective number of sequences. We did not impose any sparsity on the model. We used the profile model to initialize the parameters, that is, all fields were set to $h_i(a) = \log f_i(a) + \text{constant}$ where $f_i(a)$ is the empirical frequency and all couplings were initialized at zero. We did not impose any $L1$ and $L2$ regularization.

We used 1000 Markov chains per training epoch with 20 configurations saved per chain and used persistent chains, which means that the initial configuration for any epoch is the last configuration for the previous epoch. For sampling, we set the "wait time" $T_{wait}$ equal to 20, corresponding to 20 sweeps where 1 sweep is equal to $L = 51$ (the length of the sequence) Monte Carlo steps. The equilibration time $T_{eq}$ is then set to $2T_{wait}$. Then the $n^{th}$ configuration is sampled every $T_{eq} + nT_{wait}$ sweeps.

For outputting, adabmDCA performs one final sampling and outputs the fields and couplings in a file, along with the sampled configurations. These couplings and fields were used to calculate the Hamiltonian associated with the model in order to find the log fold improvement in the probabilities. The functions used to calculate these values can be found at github.com/hodaakl/GENERALIST/benchmark/adabmDCA.

**VAE:** For the VAE, we used the architecture given in the original manuscript [7] and changed the latent dimensions and the necessary parameters to use an MSA of different sizes. Following the procedure in the original manuscript, the sequences of the given MSA were clustered using mmseqs2 [8], using a sequence identity threshold of 70%. Clusters were then randomly chosen for the validation set until the validation set had a size of 20% of the total number of sequences. After training, we chose the best-performing latent dimension that resulted in the most accurate model for VAE according to $r_{20}$ plot SI Fig 1. When we compare the VAE-generated ensembles with other models (GENERALIST, ArDCA, adabmDCA), we choose the optimal latent dimension (the one that most accurately reproduces the $r_{20}$ summary statistics). The optimal latent dimension was 16, 4, 4, 4, and 8 for proteins BPT1, DHFR, P53, EGFR and mTOR respectively.

#### 5 Identifying model-predicted local optimal sequences

This analysis is performed on ArDCA, adabmDCA and GENERALIST models trained on protein BPT1. To find the local optimal sequences given adabmDCA and ArDCA models, we use 5000 natural sequences and mutated them to identify sequences with local optimum probability. We only considered natural sequences without alignment gaps and mutations that introduced gaps were not considered in the optimization process.

At any stage in the search process, we perform a random single mutation and assess the relative probability ratio of the proposed mutant compared to the starting sequence. If the mutant increases in probability, it is accepted, otherwise rejected. We repeated this process until the algorithm gets "stuck" on a sequence for $200L$ steps , where $L = 51$ is the length of BPT1 sequence, i.e. for $200L$ consecutive single-site mutations, no sequence is accepted.

For GENERALIST, the search for the local optima is through obtaining the maximum probability sequence corresponding to a given $z$. Those $z$s are the embeddings of the same natural sequences used as the starting sequences of the ArDCA/adabmDCA sequence search.

#### 6 Structure prediction using AlphaFold2

We predict the structures of locally optimal sequences of ArDCA and adabmDCA and GENERALIST as well the natural sequences used as the starting sequences for the local optima search algorithm using AlphaFold2 (AF2) [9]. For any ensemble (natural or generated), 5000 sequences are used. We provide the natural MSA as the context MSA for all structure predictions. The structure prediction provides a predicted LDDT (plddt) score per residue position. This value is a proxy for AF2 confidence in the predicted structure, as well as the degree of order in the sequence region. We use the average of plddt for each sequence to compare the structure quality of local optima found by different models.

Figure 1: **SI figure 1. Panel A.** Box plot of fractional Hamming distance distributions to closest natural sequence for VAE model of different latent dimensions (x labels). **Panel B.** The average Pearson correlation coefficient between frequencies of top 20 amino acid combination of order n (x-axis) averaged across different combinations (y-axis) for VAE model trained with different latent dimensions (legend). For both panels, each subplot represents a different protein from left to right: BPT1, DHFR, P53, EGFR, mTOR.

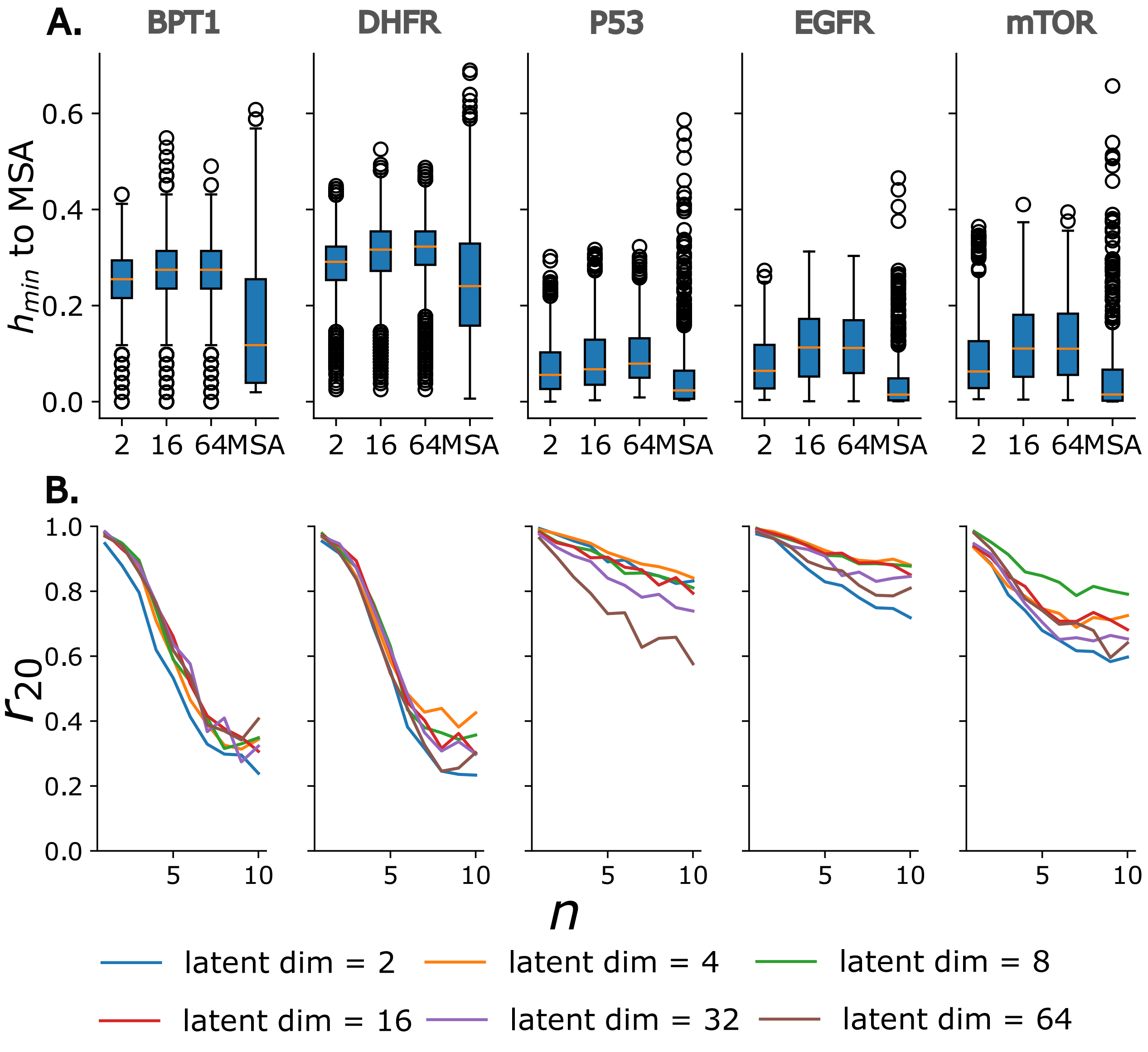

Figure 2: **SI figure 2. Panels A, B and C.** For each order of statistics (grey legend panel A), the frequencies of different amino-acid strings are obtained from Natural and generated ensemble. For order$\geq 2$, mean removed frequencies are used. The slope of the best fit line of those frequencies as well as the Pearson coefficient of correlation is obtained. 1 - slope (y-axis) vs 1 - Pearson correlation (x-axis) are plotted for different models (legend). Panel **A**, protein DHFR, panel **B** protein P53 and panel **C**z protein mTOR.

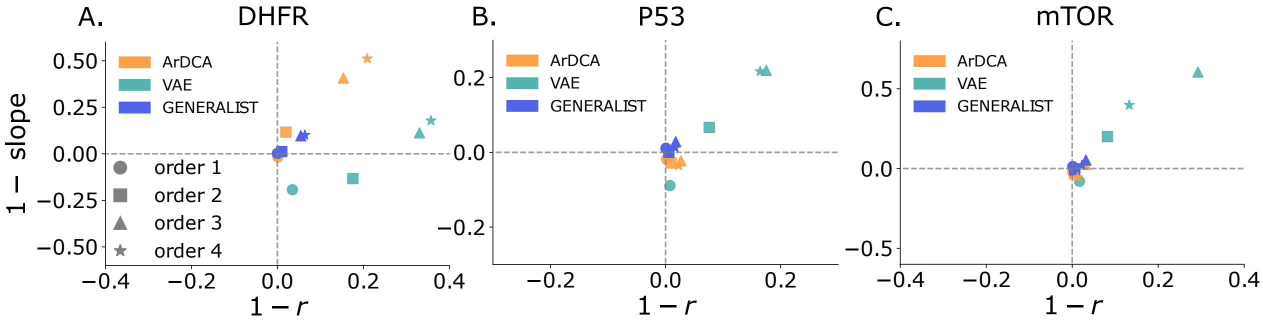

Figure 3: **SI Figure 3. Panels A, B, and C.** The average Pearson correlation coefficient between frequencies of top 20 amino acid combinations of order n (x-axis) averaged across different combinations (y-axis). Panel **A**, protein DHFR, panel **B** protein P53 and panel **C** is protein mTOR.

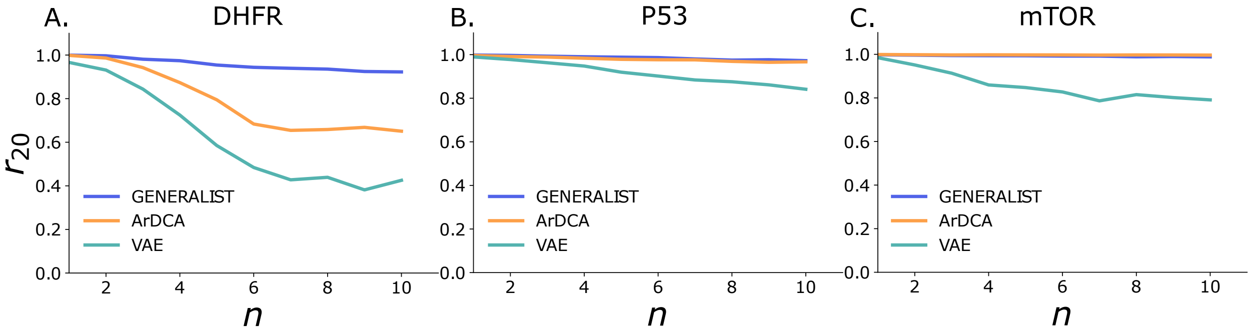

Figure 4: **SI Figure 4. Panels A, B** The average Pearson correlation coefficient between frequencies of the top 10 amino acid combinations of order n (x-axis) averaged across different combinations (y-axis). **Panels C, D** Same as A and B but using the frequencies of the top 50 amino acid combinations to obtain the average Pearson correlation. Panels **A** and **C** for protein BPT1 and panels **B** and **D** for protein EGFR

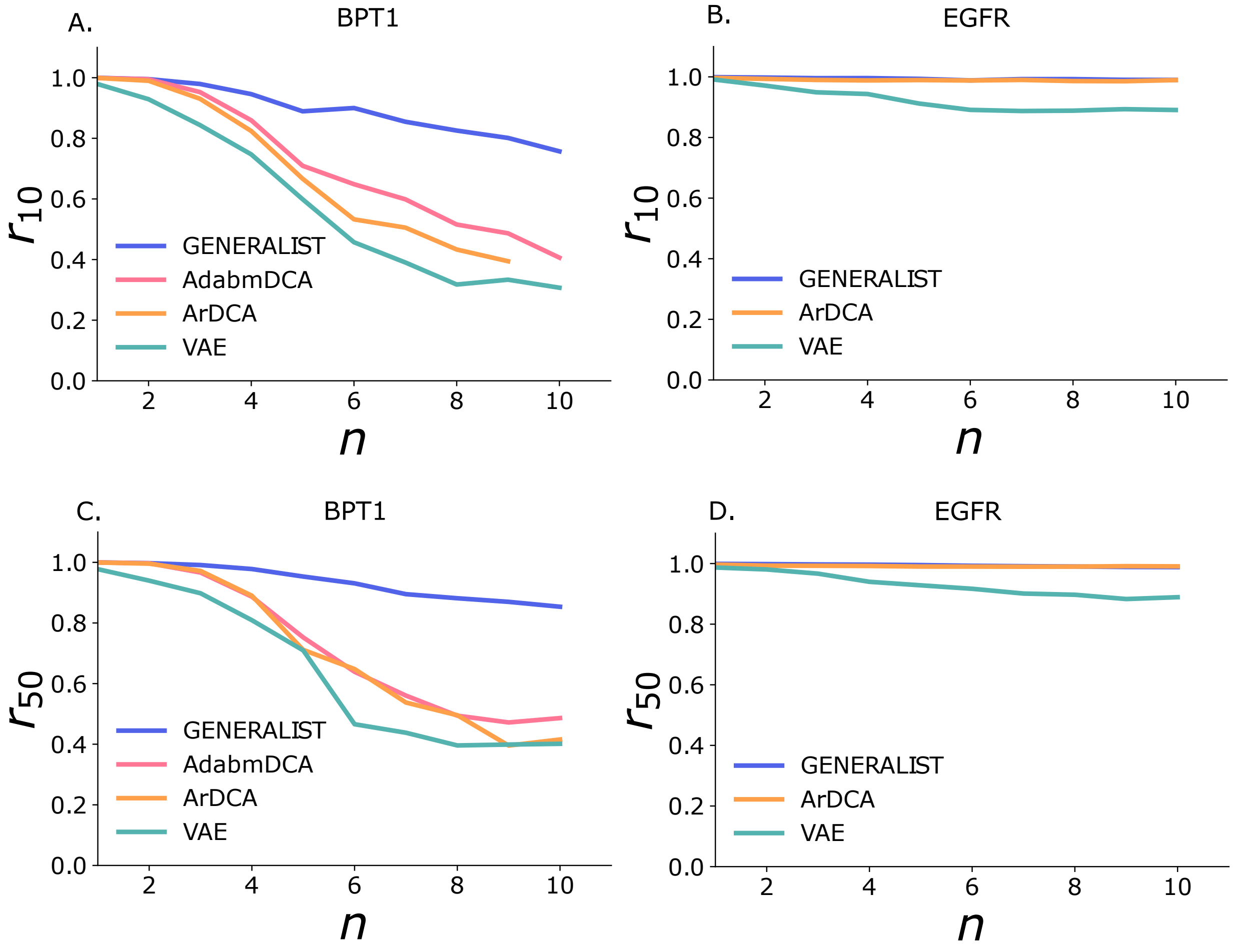

Figure 5: **SI Figure 5** Distribution of fractional Hamming distances between random pairs of sequences within an ensemble shown for DHFR (panel **A**), P53 (panel **B**) and mTOR (panel **C**).

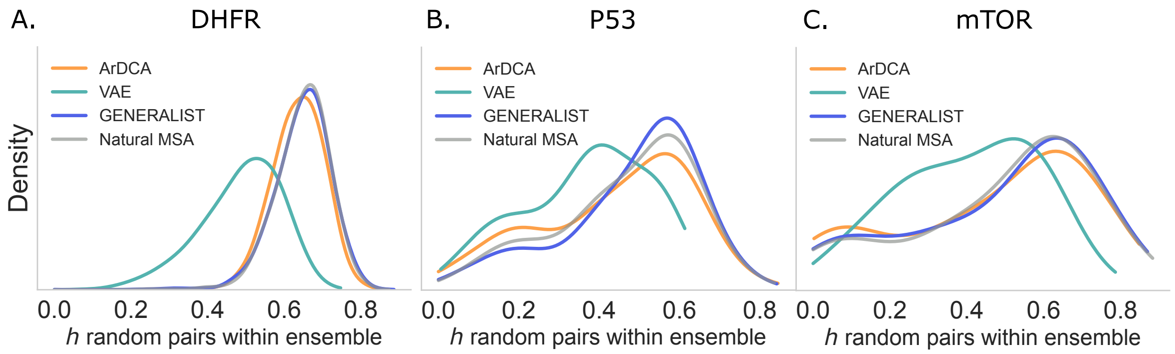

Figure 6: **SI Figure 6.** Distribution of fractional Hamming distances to the closest sequence within an ensemble for different models shown for DHFR (panel **A**), P53 (panel **B**) and mTOR (panel **C**)

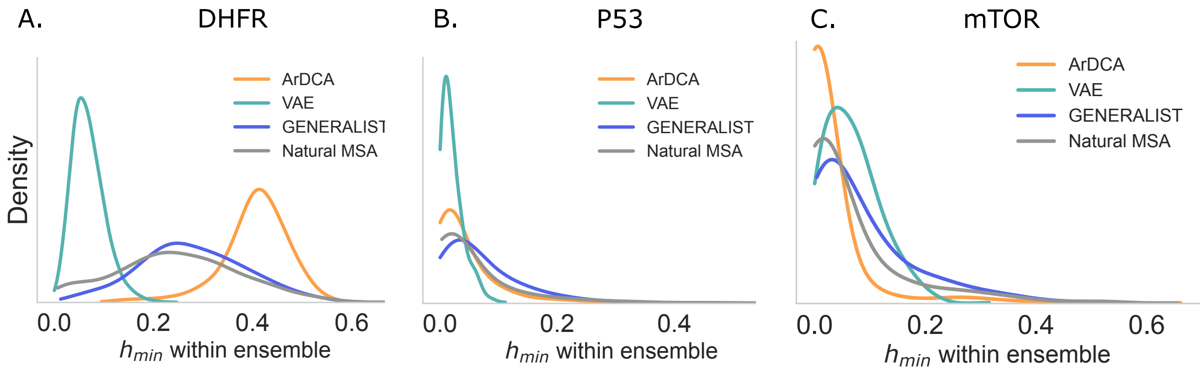

Figure 7: **SI Figure 7.** Distribution of fractional Hamming distances to closest natural sequence for different models shown for DHFR (panel A), P53 (panel B), and mTOR (panel C).

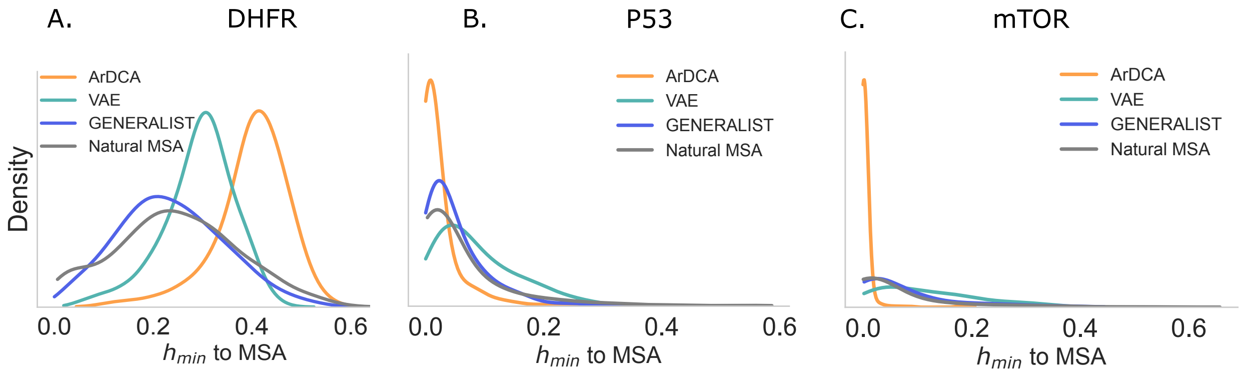

[2] Jaina Mistry, Sara Chuguransky, Lowri Williams, Matloob Qureshi, Gustavo A Salazar, Erik L L Sonnhammer, Silvio C E Tosatto, Lisanna Paladin, Shriya Raj, Lorna J Richardson, Robert D Finn, and Alex Bateman. Pfam: The protein families database in 2021. *Nucleic Acids Research*, 49(D1):D412–D419, Jan 2021.

[3] Hmmer, version 3.2.1 (june 2018). `http://hmmer.org/`.

[4] Francisco McGee, Sandro Hauri, Quentin Novinger, Slobodan Vucetic, Ronald M. Levy, Vincenzo Carnevale, and Allan Haldane. The generative capacity of probabilistic protein sequence models. 12(1):6302.

[5] Anna Paola Muntoni, Andrea Pagnani, Martin Weigt, and Francesco Zamponi. adabmdca: adaptive boltzmann machine learning for biological sequences. *BMC Bioinformatics*, 22(1):528, Dec 2021.

[6] Jeanne Trinquier, Guido Uguzzoni, Andrea Pagnani, Francesco Zamponi, and Martin Weigt. Efficient generative modeling of protein sequences using simple autoregressive models. *Nature Communications*, 12(1):5800, Dec 2021.

[7] Alex Hawkins-Hooker, Florence Depardieu, Sebastien Baur, Guillaume Couairon, Arthur Chen, and David Bikard. Generating functional protein variants with variational autoencoders. *PLOS Computational Biology*, 17(2):e1008736, Feb 2021.

[8] Martin Steinegger and Johannes Söding. Mmseqs2 enables sensitive protein sequence searching for the analysis of massive data sets. *Nature Biotechnology*, 35(11):1026–1028, Nov 2017.

[9] John Jumper, Richard Evans, Alexander Pritzel, Tim Green, Michael Figurnov, Olaf Ronneberger, Kathryn Tunyasuvunakool, Russ Bates, Augustin Žídek, Anna Potapenko, Alex Bridgland, Clemens Meyer, Simon A. A. Kohl, Andrew J. Ballard, Andrew Cowie, Bernardino Romera-Paredes, Stanislav Nikolov, Rishub Jain, Jonas Adler, Trevor Back, Stig Petersen, David Reiman, Ellen Clancy, Michal Zielinski, Martin Steinegger, Michalina Pacholska, Tamas Berghammer, Sebastian Bodenstein, David Silver, Oriol Vinyals, Andrew W. Senior, Koray Kavukcuoglu, Pushmeet Kohli, and Demis Hassabis. Highly accurate protein structure prediction with alphafold. *Nature*, 596(7873):583–589, Aug 2021.
